# Hippocampal sharp-wave ripples correlate with naturally occurring self-generated thoughts in humans

**DOI:** 10.1101/2023.06.23.546219

**Authors:** Takamitsu Iwata, Takufumi Yanagisawa, Yuji Ikegaya, Jonathan Smallwood, Ryohei Fukuma, Satoru Oshino, Naoki Tani, Hui Ming Khoo, Haruhiko Kishima

## Abstract

Core features of human cognition, for example, the experience of mind wandering, highlight the importance of the capacity to focus on information separate from the here and now. However, the brain mechanisms that underpin these self-generated states remain unclear. An emerging hypothesis is that self-generated states depend on the process of memory replay, which, in animals, is linked to sharp-wave ripples (SWRs) originating in the hippocampus. SWRs are transient high-frequency oscillations that exhibit circadian fluctuations and, in the laboratory, are important for memory and planning. Local field potentials were recorded from the hippocampus of 11 patients with epilepsy for up to 15 days, and experience sampling was used to describe their association with ongoing thought patterns. SWRs were correlated with patterns of vivid, intrusive ongoing thoughts unrelated to the task being performed, establishing their contribution to the ongoing thoughts that humans experience in daily life.

## Introduction

Both in laboratory settings and in daily life, human cognition often escapes the here and now to focus on information unrelated to the immediate environment^1,2^. Estimates suggest that in daily life, the human mind can wander from the immediate environment for up to 30% of the time^2^, with patterns of self-focused future thought characteristically occurring during activities such as exercising or commuting, while patterns of distracting intrusive thoughts occur frequently at rest^3^. An emerging body of literature has begun to uncover the complex associations of these states with intelligence^4–6^, autism^7^, attention-deficit disorder^8^ and happiness and well-being^2,9^. Neuroimaging evidence suggests that these processes are linked to activity in regions such as the posterior cingulate and the medial prefrontal cortex, both of which are key nodes in what is known as the default mode network^10–12^. Likewise, studies examining variation in cortical anatomy have highlighted individual differences in the medial temporal lobe as important for trait variation in these experiences in both the laboratory and daily life^13,14^.

Despite this emerging literature, we know relatively little about how these experiences are orchestrated. Contemporary views on these experiences hypothesize that these experiences may be linked to hippocampal function^15–17^ given its role in memory and prospection^18^ as well as the possibility that the medial temporal lobe plays a general role in the organization of cortical function^19,20^. One possibility is that the process of self-generated thought is linked to the emergence of sharp-wave ripples (SWRs) in the hippocampus. SWRs are bursts of synchronized neuronal activity in the mammalian hippocampus, which vary depending on the state of the animal^21,22^. SWRs frequently occur during non-REM sleep associated with memory consolidation^21–23^; during wakefulness^24,25^, SWRs increase when animals stay in a quiescent motionless state and decrease during active movements^21,22^, a situation that—in humans—is linked to the emergence of intrusive thoughts^3^. Moreover, SWRs during wakefulness are believed to aid in memory retrieval and guide decision making based on past experiences^26,27^. Recent studies using human intracranial recordings have revealed that SWR rates in humans are related to mental contents that emerge in episodic recollection during cognitive tasks^23,28^. In light of these observations coupled with evidence that lesions to the hippocampus can decrease experiences such as mind wandering^29^, our study explored the hypothesis that SWR fluctuations are linked not only to circadian rhythms and physical activities ^21,22^ but also to accompanying mental states that are less closely coupled to the immediate environment.

In our study, we continuously measured human hippocampal local field potentials (LFPs) for 9 to 15 days in 11 patients with intracranial electrodes implanted in or adjacent to the hippocampus for the treatment of refractory epilepsy (Supplementary Table 1). All patients were monitored by video camera while they freely engaged in normal activities during neural activity recording (Fig. 1). In parallel with the LFP recordings, we assessed the patients’ mood and thought contents using the modified Differential Emotions Scale (mDES)^30,31^. Specifically, patients were asked to rate their experiences along 17 dimensions using a tablet device once every hour (see Table 1 for the list of dimensions). Additionally, we simultaneously recorded information on the patients’ physical states, such as interbeat interval (IBI), electrodermal activity (EDA), accelerometer (ACC), and blood volume pulse (BVP) data, using a wearable device attached to their left arm. We analyzed the associations between these simultaneously recorded signals to better understand how human SWRs change over the circadian rhythm and relate not only to individuals’ physical states but also to the content of their thoughts.

**Table 1.** Multidimensional experience sampling questionnaire on thoughts and feelings. For each question, subjects were asked to “rate how you feel or what you are thinking about right now” on a scale from 1 to 7 by selecting one of seven buttons representing internal states. All questions were asked using a tablet.

| Dimension | Questions | 1 | 7 |
| --- | --- | --- | --- |
| External-task Focus | My thoughts were focused on the task I was performing. | Not at all | Completely |
| Future | My thoughts involved future events. | Not at all | Completely |
| Past | My thoughts involved past events. | Not at all | Completely |
| Self | My thoughts involved myself. | Not at all | Completely |
| Person | My thoughts involved other people. | Not at all | Completely |
| Emotion | The content of my thoughts was: | Negative | Positive |
| Images | My thoughts were in the form of images. | Not at all | Completely |
| Words | My thoughts were in the form of words. | Not at all | Completely |
| Vivid | My thoughts were vivid as if I was there. | Not at all | Completely |
| Detail | My thoughts were detailed and specific. | Not at all | Completely |
| Habit | This thought has recurrent themes similar to those I have had before. | Not at all | Completely |
| Unwilling | My thoughts were what I did not want to think. | Not at all | Completely |
| Evolving | My thoughts tended to evolve in a series of steps. | Not at all | Completely |
| Spontaneous | My thoughts were: | Spontaneous | Deliberate |
| Happy | My feeling was: | Sad | Happy |
| Calm | My feeling was: | Restless | Calm |
| Excited | My feeling was: | Boring | Excited |

**Fig. 1.**
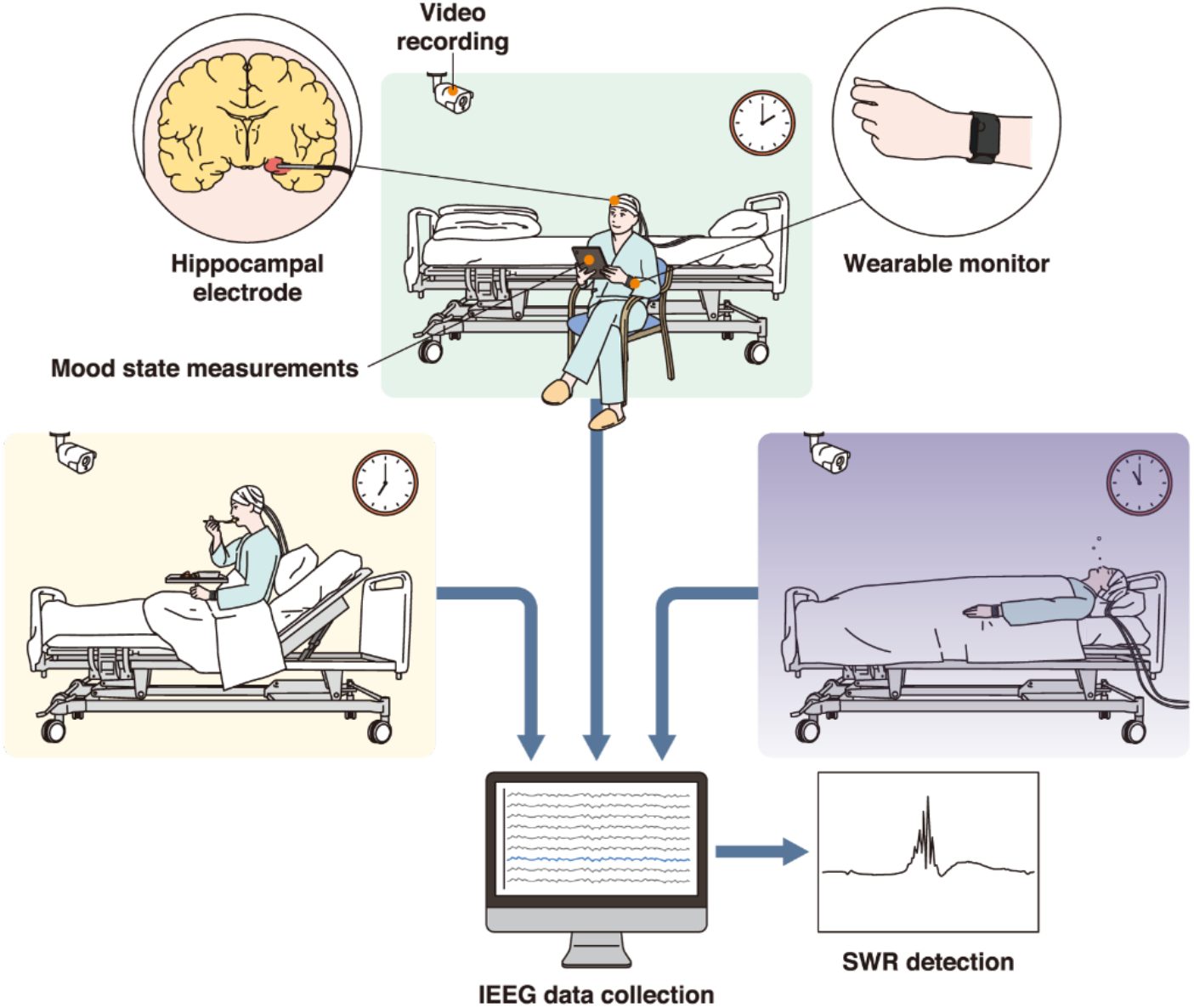
Experimental environment. We continuously measured human hippocampal LFPs for 9 to 15 days in 11 patients who had intracranial electrodes implanted in or adjacent to their hippocampus for the treatment of refractory epilepsy. All patients were monitored by two video cameras in a room where they freely engaged in normal activities during LFP recording. In parallel with the LFP recordings, we assessed the patients’ moods and thoughts using a tablet once every hour. Additionally, we simultaneously recorded the patients’ physical state using a wearable device attached to their left arm.

## Results

### Diurnal fluctuation in hippocampal SWR rates in freely behaving humans

SWRs were assessed using LFP recordings starting 4 days after electrode implantation.

We evaluated the LFP signals between two electrodes located either within or near the hippocampus. The electrode locations were identified using pre- and postoperative computed tomography (CT) and magnetic resonance imaging (MRI) scans (Fig. 2a). The periods associated with epileptic activity or movement artifacts were excluded to detect ripple candidates. The LFP of hippocampal electrodes in the selected site was then converted to a bipolar signal. After removing the power-line noise, the signals were filtered between 70 and 180 Hz to select ripple candidates. Ripple candidates whose durations were shorter than 20 ms or longer than 200 ms were rejected, and peaks less than 30 ms apart were concatenated. The number of detected SWRs during each minute was *Z*-standardized over each day to obtain an SWR rate for each subject.

**Fig. 2.**
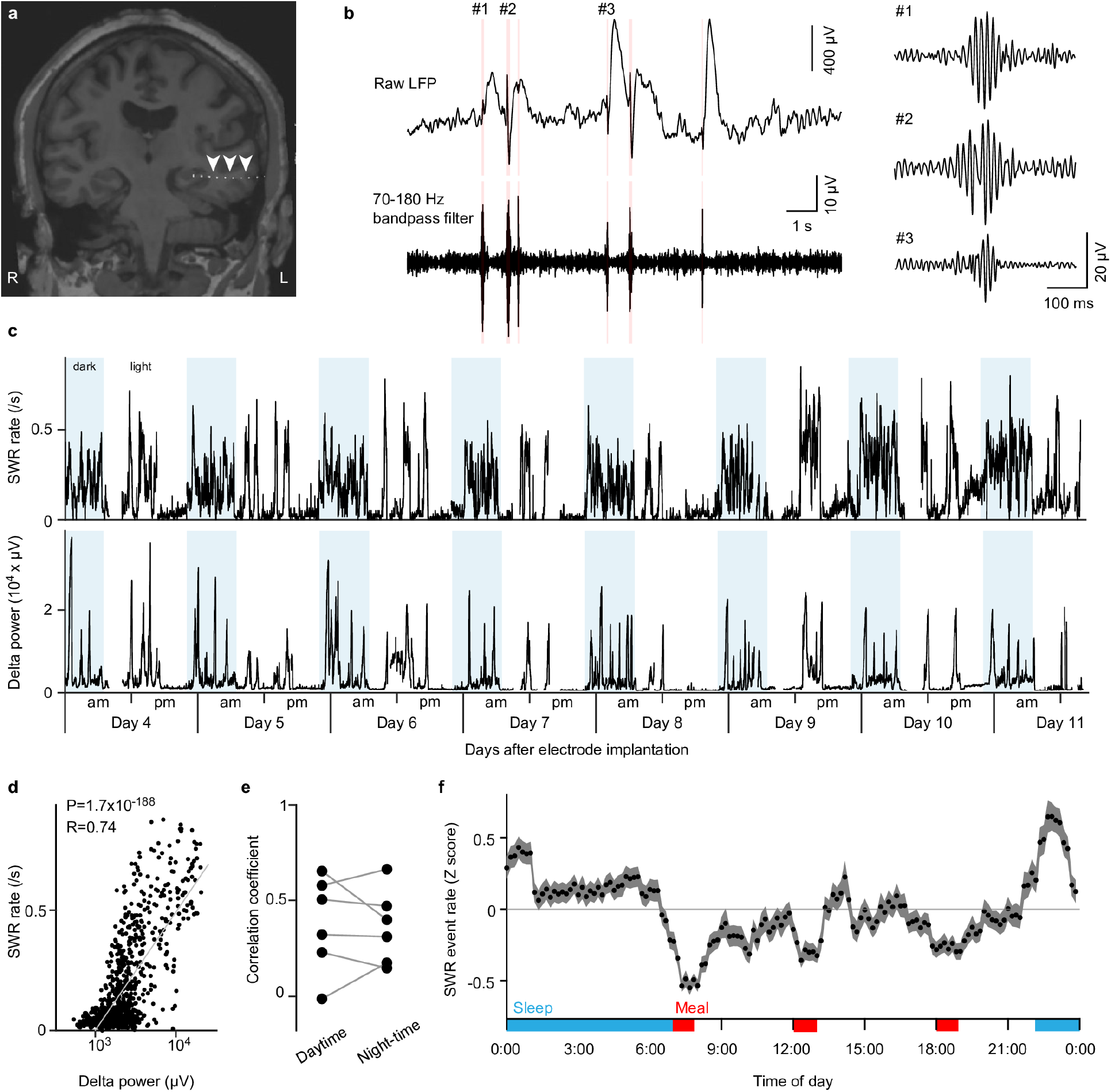
Diurnal variation in hippocampal SWR rates in freely behaving humans. A, An image combining preoperative MRI and postimplantation CT data illustrates the placement of electrodes within the hippocampus. B, Hippocampal LFPs (top) were bandpass-filtered between 70 and 180 Hz (bottom). Three representative SWRs (asterisks) are magnified in the right inset. C, A time course of SWR rates (top) and neocortical delta power (bottom) for 8 consecutive days (top) in a representative patient. The light blue background indicates the time of day when theroom was darkened. D, SWR rates are positively correlated with delta power as shown in Panel c. Each dot represents a single time bin of 10 min. e, The correlation coefficients between SWR rates and delta power did not differ between daytime and nighttime in a total of six patients with implanted cortical electrodes. F, The *Z* score mean and 95% confidence interval of the diurnal variation in SWR rates over 24 hr in all 11 patients. The red and blue bars indicate meal and sleep times, respectively.

Fig. 2c (top) shows the SWRs of a representative patient (Pt-04); overall SWR rates increased during the night, exhibiting intense rate fluctuations, and decreased upon waking. In six patients with implanted cortical and hippocampal electrodes, we computed the delta wave amplitude, an indicator of slow-wave sleep, from the neocortical electrodes to examine their relationship with SWR rates (Fig. 2c bottom). Over seven days of recording, SWR rates increased when delta amplitudes increased (Fig. 2d; *R* = 0.74, *P* = 1.69× 10^−188^, *n* = 1,296 time points, Pearson correlation coefficient). The correlations were similar during the day and night (Fig. 2e; *R* = 0.38 ± 0.25 for daytime and 0.36 ± 0.19 for nighttime; *t*5 = 0.2993; *P* =0.7768; paired *t* test). These data are consistent with a previous report demonstrating that SWR rates vary with sleep cycles.^21^ For subsequent analysis, the SWR rates were *Z*-standardized for each day, averaged for each patient, and pooled among all eleven subjects. The data indicated a significant increase in SWR rates during sleep and a decrease during the daytime (Fig. 2f; *F*143,94391 = 42.1, *n* =126,720 time points from a total of 88 days in 11 patients; one-way ANOVA). Another salient feature of SWR rates consisted of transient drops at 7:00, 12:00, and 18:00, coinciding with the scheduled mealtimes in the hospital, with the rates returning to baseline levels after meal completion. During the day, the SWR rates were higher during the afternoon than during the morning (morning, -0.24 ± 0.87; afternoon, -0.11 ± 0.94; *P* < 0.001, *t*58,384 = -16.5, *n* =83,700 time points, *t* test). These findings indicate that SWR rates exhibit characteristic circadian rhythms in humans under freely behaving conditions.

### Correlation between SWR rates and biophysical parameters

We examined whether the SWR rates were associated with changes in the activity and autonomic state of patients. Specifically, the EDA, ACC, BVP, and IBI of the patients were *Z*-standardized and averaged across patients (Fig. 3a). We then used a linear regression analysis to predict the SWR rates based on the four measurements and time of day (hour). The predicted SWR rates were weakly but significantly correlated with the actual SWR rates (*R* = 0.21, *P =* 4.3 × 10^−10^, *n* = 2,027 time points) (Fig. 3b). Among the five factors examined, BVP (*R* = -0.007, *P* = 6.6 × 10^−3^) and IBI (*R* = 0.15, *P* = 4.4 × 10^−11^) significantly explained diurnal variations in SWR rates as a function of time of day (Fig. 3c).

**Fig. 3.**
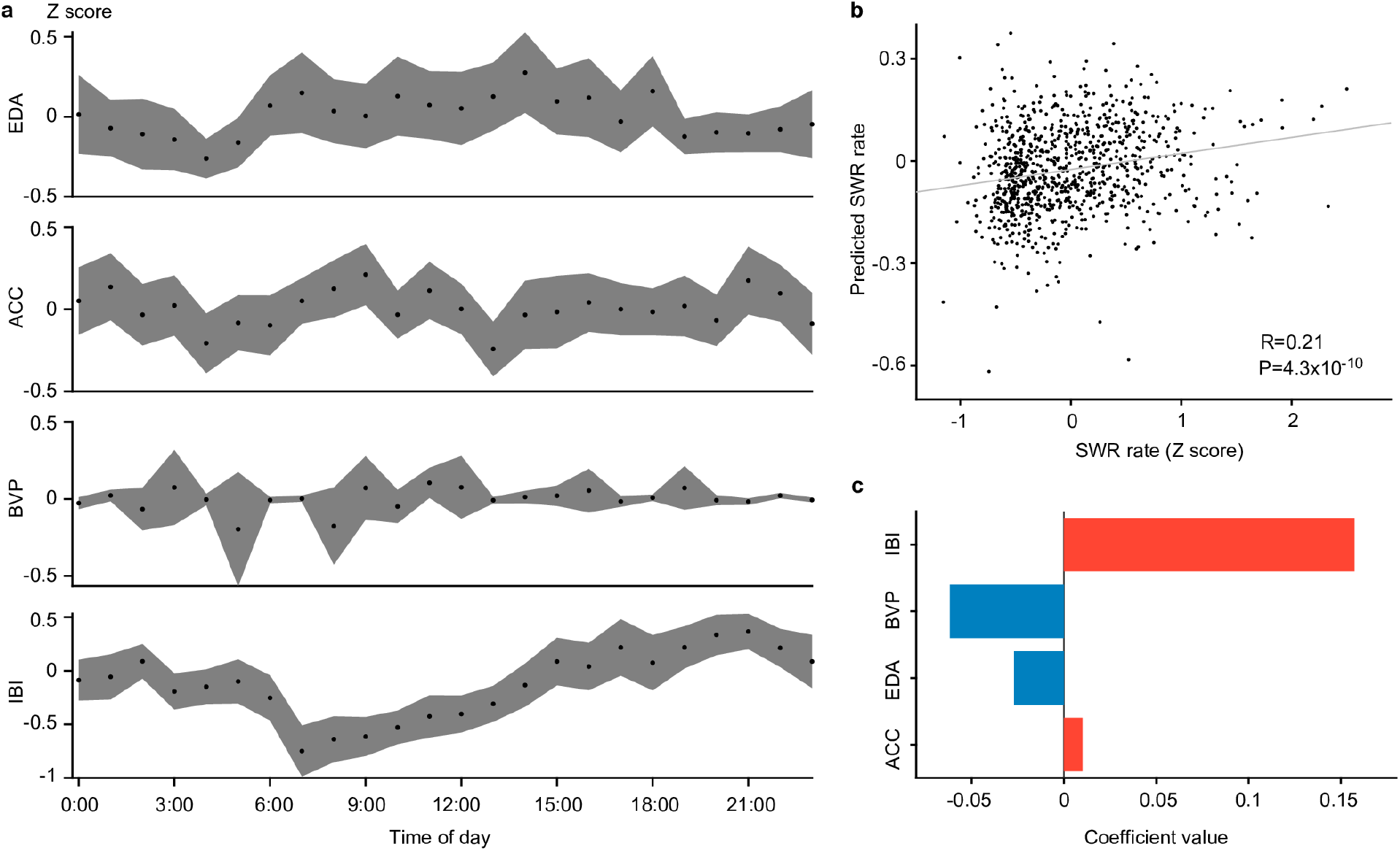
Correlation between SWR rates and biophysical parameters. A, Diurnal variations in the mean and 95% confidence of Z scores of EDA, ACC, BVP, and IBI values of all 11 patients. B, A linear regression model was used to predict SWR rates from EDA, ACC, BVP, and IBI values. Each dot represents a single prediction for a 60-min period. C, Coefficient of each predictor in the regression model. The red and blue bars indicate negative and positive coefficients, respectively.

### Correlations of SWR rates with thoughts and feelings

To investigate the correlation between SWR rates and thought patterns, the patients answered mDES questions every hour between 8 am and 8 pm. They were asked to indicate their thought content (14 questions) and mood (3 questions) on a Likert scale from 1 to 7. Before the data were analyzed, the responses were standardized for each participant. The analysis revealed that only the spontaneity of cognition significantly varied throughout the day, with the highest levels observed in the mid-afternoon and the lowest levels in the early evening (*P* = 0.033, *F*13,267 = 1.88; one-way ANOVA) (Fig. 4a). We used a linear regression model to predict the average SWR rates in the 2 to 7 min preceding each questionnaire response based on the standardized values of the responses to each question. The predicted values from the regression model showed a significant correlation with the actual SWR rates (*R* = 0.47, *P* =1.7 × 10^−10^, *n* = 192 questions) (Fig. 4b), which was significantly stronger than the correlation observed between SWRs and measures of active and autonomic responses (*Z* = 0.51 versus 0.21, *P* = 6.9 × 10^−5^, Fisher’s *Z* test), highlighting specific correlations of SWRs with patterns of ongoing thought. Specifically, we found that across the 17 dimensions analyzed, the patients reported less external-task focus (estimated coefficient (EC) = -0.075, *P* =2.2 × 10^−5^), less desired experiences (EC = 0.049, *P* = 6.4 × 10^−3^), and more vivid mental content (EC = 0.036, *P* = 0.0219) the greater the SWR rate (Fig. 4c). However, we did not observe any significant correlation between mood state and fluctuations in SWR rates. These findings establish that SWRs are specifically related to patterns of ongoing thoughts that humans experience in daily life.

**Fig. 4.**
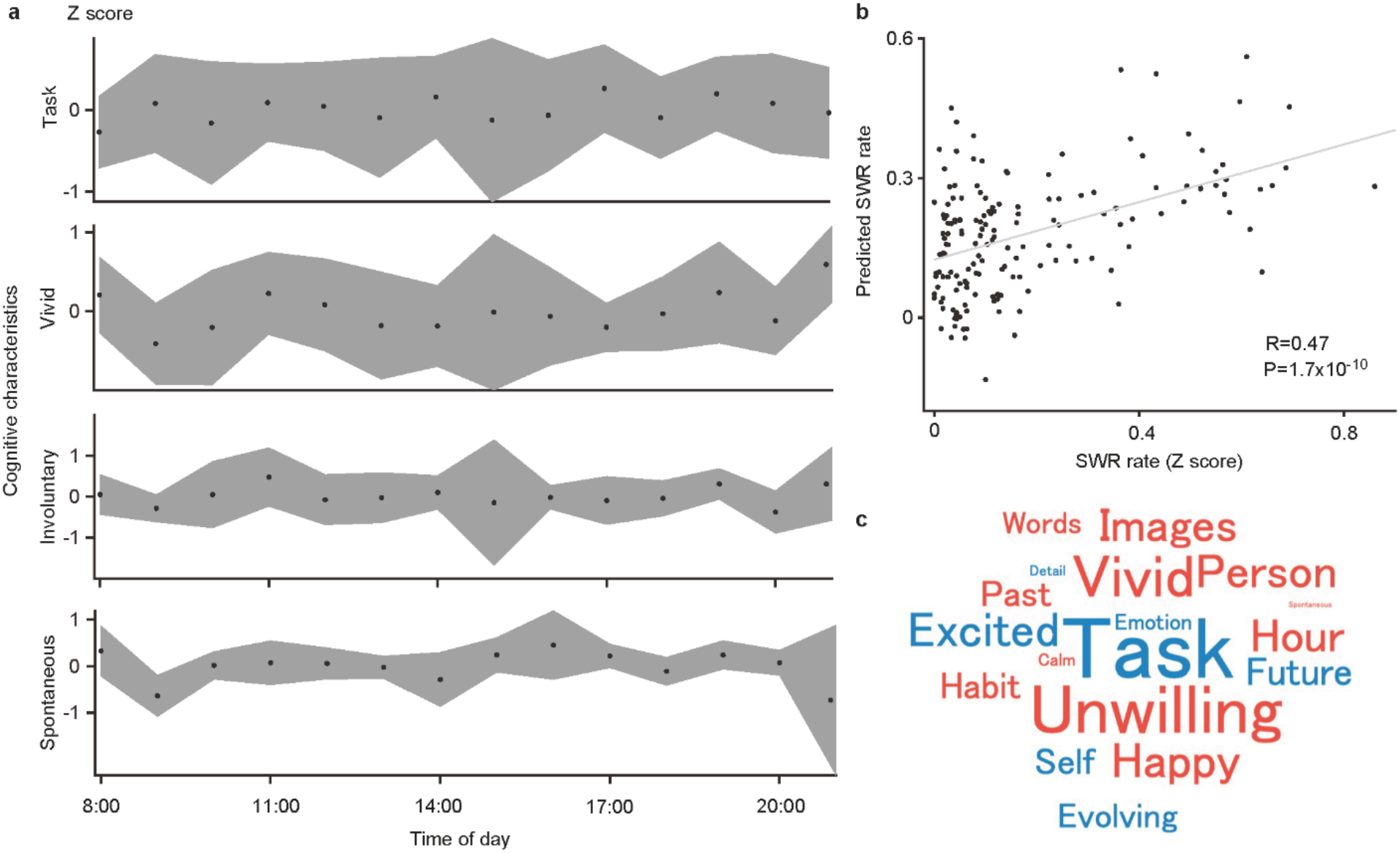
Correlations of SWR rates with thoughts and feelings. A, Diurnal variations in the mean and 95% confidence of Z-score data on focused, vivid, involuntary, and spontaneous thoughts in all 11 patients. B, A linear regression model was used to predict SWR rates from 17 features. Each dot represents a single prediction for a 5-min period. C, A word cloud displaying the coefficient of each feature in the linear regression model. Red and blue indicate positive and negative coefficients, respectively, and the size of each word indicates the magnitude of the absolute value of the coefficient. *Note that only the spontaneous nature of cognition varied with time of day; the other three dimensions of experience are included in Panel A because they were related to the occurrence of hippocampal SWR*.

## Discussion

Hippocampal SWRs have been extensively studied in relation to memory processes; however, their relationship to naturally occurring patterns of ongoing thoughts in humans is not yet fully understood. Here, we report the first evidence that human SWRs are related to the content of ongoing thoughts in a manner that is consistent with an emerging hypothesis linking the medial temporal lobe to patterns of self-generated thought^15^. Furthermore, our findings demonstrated a diurnal fluctuation in freely behaving humans, characterized by a decrease in SWR activity around mealtimes and an increase in the afternoon compared to the morning. Previous studies in rodents demonstrated that SWRs occur during nonexploratory states, such as slow-wave sleep, rest, grooming, and eating/drinking^21^. As blood glucose concentrations are inversely correlated with SWR rates^32^, the hippocampus may help regulate hormones and the autonomic nervous system. Furthermore, the positive correlation between SWR rates and IBI suggests that the frequency of SWRs is increased in a parasympathetic-dominant state, which is consistent with the increased occurrence of SWRs during non-REM (NREM) sleep^21^.

Interestingly, in our study, SWRs were more strongly associated with the content of ongoing thoughts than with physical activity.

Animal studies are limited in terms of their ability to investigate cognition without direct links to behavior; however, using experience sampling, we established that SWR rates were correlated with descriptions of vivid, unwanted ongoing thoughts that were unrelated to the current task. This finding supports an emerging hypothesis implicating the medial temporal lobe in forms of self-generated cognition. For example, hippocampal lesions are associated with reductions in off-task thought in the form of mind wandering^29^. In healthy adults, reports of ongoing thoughts with more vivid features are linked to greater gray matter volumes in the posterior parahippocampus, while volume in the anterior parahippocampus is linked to off-task states^13^. Furthermore, the hippocampus is functionally aligned with the default-mode network^28,33^, and functional MRI studies show that self-generated contents are linked to activity in anterior regions of this system, while vivid and detailed experiences have links to posterior regions^20,34,35^. Finally, SWRs are also important in laboratory tasks that mimic naturally occurring self-generated thought, including future planning, memory recall, and imagination^23,36^—situations that depend on the self-generation of mental content and can also include a reduction in the perceptual processing of external input^37^. Together with this evidence, therefore, our study suggests that SWR in the hippocampus contributes to naturally occurring ongoing thought patterns in humans.

Our study also has important clinical implications because it links SWRs in the hippocampus to unwanted ongoing thoughts, a type of thinking that is common in daily life^38^ and that is prevalent in conditions such as obsessive-compulsive disorder^38^, anxiety and depression^39^ and posttraumatic stress disorder^40^. Our evidence highlighting associations with unwanted thoughts and the hippocampus and ongoing thoughts using experience sampling will provide a method for elaborating on contemporary views of intrusive thoughts as emerging when the hippocampus is no longer regulated by the prefrontal cortex^41^.

Although our data establish a correlation between SWRs in the hippocampus and the content of ongoing thought patterns, there are several limitations that should be borne in mind when considering these results. First, while we carefully excluded the effects of epileptic activity, it remains unclear whether epileptic activity influenced hippocampal function or led to changes in ongoing patterns of thought. Although intracranial recordings are the gold standard for linking hippocampal activity to cognition^16^, it will be important for future studies to use high-field imaging methods^42^ to establish how neural activity within the medial temporal lobe contributes to ongoing thoughts not only in patients but also, critically, in healthy controls.

Second, the recordings were conducted in a hospital, rather than in the patients’ homes or other natural environments. This situation constrains the possible activities patients can engage in and consequently may impact the accompanying patterns of thoughts^3^. It is possible, therefore, that SWR contributes to the features of ongoing thoughts more broadly than was observed in our study. Third, our data are correlational; thus, further studies that manipulate SWRs and associated cognitive functions are needed to fully understand the role the hippocampus plays in naturally occurring features of human cognition. Since SWRs in the hippocampus are influenced by neuromodulators, it is possible that pharmacological manipulation of these systems may provide the opportunity to test causal accounts of the role that SWRs play in ongoing thought patterns in humans^17^.

In conclusion, our analysis of long-term intracranial electroencephalography (EEG) data from the human hippocampus has demonstrated a diurnal fluctuation in hippocampal SWRs during periods of wakefulness. Our findings suggest that these SWRs not only change over the sleep-wake cycle but also vary in association with the emergence of human thought contents that are unrelated to the immediate environment. Our study paves the way for future studies of continuous intracranial EEG and thought sampling that will help clarify the function that SWRs play in human cognition.

## Methods

### Participants

In our hospital, 23 patients with drug-resistant epilepsy underwent intracranial electrode implantation to treat epilepsy between January 2020 and May 2022. Of these, 15 patients (16 electrodes) had deep electrodes implanted in the hippocampus or parahippocampal gyrus. To minimize the influence of epileptic activity, recordings were excluded from the analysis if the hippocampus was pathologically diagnosed with hippocampal sclerosis^16^. Ultimately, eleven patients with eleven electrodes (6 males; age: 32.8 ± 13.8 years; mean ± standard deviation [SD]; see Supplementary Table 1) who consented to participate in this study (i.e., to complete the questionnaire and wear a wearable device) were included. Electrode placement was determined solely by clinical necessity. In all patients, the clinical team determined the placement of the electrodes to best localize epileptogenic regions. The EEG data analyst did not participate in the determination of electrode placement. Data were collected at the Department of Neurosurgery at Osaka University Hospital. The research protocol was approved by the Institutional Review Board (approval no. 14353, UMIN000017900), and informed consent was obtained from the participants. All data were acquired in the participants’ rooms during their hospital stays. The analysis included only EEG data from four days post implantation onward to minimize the impact of surgery. Based on visual inspection of the recorded EEG data, seizures and segments of intense epileptic activity were discarded from subsequent analyses.

### Behavioral data

Postoperatively, patients were able to walk around in the hospital room with the intracranial electrodes connected to the EEG system (Fig. 1). The connection to the EEG system was temporarily disconnected for bathing or clinical examinations. Within the room, patients were constantly recorded by video synchronized with EEG data. The initiation and termination of eating and sleeping were determined based on the simultaneously recorded video.

#### Physiological signals

We simultaneously recorded subjects’ physiological states, such as EDA, three-dimensional acceleration data, BVP, and IBI, with the Empatica E4 wristband (Empatica, Milan, Italy) worn on their left arm. Physiological data were stored within the tablet that presented the questionnaire and synchronized with the EEG data via a time-to-live (TTL) signal generated at the time of response. Data from the EDA sensor are expressed in microsiemens, and the sampling rate was 4 Hz. Acceleration data were measured with a 3-axis accelerometer with a sampling rate of 32 Hz in units of 1/64 × *g* (gravitational acceleration); the RMS of acceleration was calculated on the 3 axes as ACC. BVP data were obtained through photoplethysmography, and the sampling rate was 64 Hz^43^. IBI data were extracted from the BVP signal and expressed in seconds^44^. All physiological data were averaged in 1-min bins and then normalized for each day. One patient who wore the wearable device for under 2 days during the measurement period was not included in the analysis of behavioral data.

#### Questionnaire

Subjects’ mood and thought contents were measured by a tablet-based self-report questionnaire composed of 17 questions that evaluated their thought contents (14 questions) and mood (3 questions) (Table 1). The questionnaire was devised primarily to assess instances of mind wandering. The queries used to evaluate mind wandering were adapted from our previous report^45^. In each question, the subject was asked to “rate how you feel or what you are thinking about right now” (Table 1) on a continuum of internal states between 1 and 7. All questions appeared on the display hourly between 8:00 a.m. and 8:00 p.m. Patients were permitted to answer questions at any time. The TTL signal was recorded on the EEG data at the moment the participant provided the response, and that moment was considered the response time.

### Intracranial EEG

#### Data processing

We recorded intracranial EEG data from subdural electrodes and depth electrodes sampled at 10 kHz with an EEG-1200 (Nihon Kohden, Tokyo, Japan). A low-pass filter with a cutoff frequency of 3,000 Hz and a high-pass filter with a cutoff frequency of 0.016 Hz were employed for signal preprocessing, and downsampling to 2 kHz was performed using an 8^th^-order Chebyshev Type I low-pass filter before resampling. Subdural contacts were arranged in both grid and strip configurations with intercontact spacing of 10 mm and a contact diameter of 3 mm. The intercontact spacing of the deep electrode contacts ranged from 5-15 mm, with a width of 1 mm. Electrode localization was accomplished by coregistering the postoperative CT data with the preoperative T1-weighted MRI data using SYNAPSE VINCENT (Fujifilm, Tokyo, Japan). The scanners used to obtain the MRI data were the SIGNA Architect (GE Healthcare) for five subjects and the Ingenia (Philips Healthcare, Amsterdam, Netherlands) for six subjects. The CT images were obtained using the Discovery CT750 HD (GE Healthcare) for four subjects and the Aquilion Precision (Toshiba Medical Systems) for seven subjects.

#### Calculation of delta band power

The amplitudes of delta waves were determined from electrocorticography data obtained from individuals with implanted cortical electrodes to analyze the correlation between sleep status and SWRs. Bipolar potentials were obtained by utilizing all available pairs of adjacent cortical electrodes. The power-line noise at 60, 120, 180, 240 (±1.5) Hz was removed with a 1,150-order FIR band stop filter. These potentials were filtered using a low-cut filter with a frequency of 0.5 Hz and a high-cut filter with a frequency of 4 Hz, and a Hilbert transform was applied. The Hilbert function was utilized to obtain the absolute value of the instantaneous amplitude of the analyzed signal. These computations were carried out using MATLAB 2017b.

#### SWR detection

The detection method followed the method described by Norman et al.^23^. The LFP of hippocampal electrodes in the selected site was then converted to a bipolar signal. To remove power-line noise at 60, 120, 180, and 240 (± 1.5) Hz, we used a 1,150-order finite impulse response (FIR) bandpass filter. Next, the signals were filtered between 70 and 180 Hz using a zero-lag linear-phase Hamming-window FIR filter with a transition bandwidth of 5 Hz. The instantaneous amplitude was computed using the Hilbert transformation and clipped at 4 × the standard deviation (SD). Finally, the clipped signal was squared and passed through a low-pass filter before having the mean and SD calculated. Ripple candidates were selected from the range when the unclipped signal exceeded 4 × SD. Ripple candidates whose durations were shorter than 20 ms or longer than 200 ms were rejected, and peaks closer than 30 ms were concatenated (Fig. 1b). We manually reviewed and excluded data during epileptic activity and movement artifacts to confirm the SWR. The detected SWRs were normalized by the mean and SD on each day.

### Statistical analysis

Unless otherwise specified, significant differences were assessed using one-way ANOVA, and Pearson correlation analysis was used to evaluate correlations. This study used observational data, and data collection and analysis were performed using MATLAB 2017b. To conduct the regression analysis, we predetermined the total number of answers on questionnaires as more than 170 samples, which is equivalent to 10 times the number of questionnaire items, totaling 17. The data gathered for each patient were influenced by their personalized treatment plan, and the duration each participant was prepared to voluntarily contribute to the study.

#### Linear regression

Regression models were constructed to predict the occurrence of SWR events based on biometric data and answers to the self-report questionnaire. The initial model employed a linear regression analysis to predict the SWR rate using EDA, ACC, BVP, and IBI data obtained from the wearable device. The second model employed a linear regression approach to predict the mean SWR rate over a period of 2-7 min preceding the response time, as inferred from the answers to the self-report questionnaire. The regression models were evaluated through a randomized 10-fold cross-validation procedure, with correlation coefficients computed between predicted and observed values. In the cross-validation, all answers to questionnaires were randomly divided into 10 groups for training and testing. The average coefficient across all folds was calculated.

## Data availability

The datasets generated and analyzed during the current study are available on Figshare at https://figshare.com/s/568e9da3493111XXXXXX.

## Code availability

The code written to detect SWRs is available at https://figshare.com/s/84409845359e7eXXXXXX.

## Acknowledgments

We thank Dr. Satoru Saito for his help in preparing the questionnaire. This research was conducted under the Japan Science and Technology Agency (JST) Exploratory Research for Advanced Technology (JPMJER1801). This research was also supported in part by the Core Research for Evolutional Science and Technology (JPMJCR18A5) and Moonshot R&D (JPMJMS2012), Grants-in-Aid for Scientific Research from KAKENHI (20H05705), and grants from the Japan Agency for Medical Research and Development (AMED) (19dm0207070h0001, 19dm0307103 and 19dm0307008). JS was supported by a Discovery Grant from the National Engineering and Science Council of Canada, as well as an award from the New Frontiers in Research Fund.

## Author contributions

T.I. contributed to study design, data collection, data coding, data analyses and manuscript writing.

T.Y. contributed to study design, data collection, data analyses and manuscript writing and editing.

Y.I. contributed to data encoding, manuscript editing, and study design. J.S. contributed to study design, data encoding, manuscript editing, and study design. R.F. contributed to data coding, data analyses. S.O., N.T. and H.K. contributed to data collection. H.K. contributed to data collection and qualitative data analyses.

## Competing interests

The authors declare no competing interests.

## Notes

### Competing Interest Statement

The authors have declared no competing interest.

